# Veni, vidi, vici? Future spread and ecological impacts of a rapidly-expanding invasive predator population

**DOI:** 10.1101/2023.07.30.551166

**Authors:** Nelsen R. David, Corbit G. Aaron, Chuang Angela, Deitsch F. John, Sitvarin I. Michael, Coyle R. David

## Abstract

Economic and ecological consequences of invasive species make biological invasions an influential driver of global change. Monitoring the spread and impacts of non-native species is essential, but often difficult, especially during the initial stages of invasion. The Joro spider, *Trichonephila clavata* (L. Koch, 1878, Araneae: Araneidae), is a large-bodied orb weaver native to Asia, likely introduced to northern Georgia, U.S. around 2010. We investigated the nascent invasion of *T. clavata* by constructing species distribution models (SDMs) from crowdsourced data to compare the climate *T. clavata* experiences in its native range to its introduced range. We found evidence that the climate of *T. clavata*’s native range differs significantly from its introduced range and that the most similar climate in North America to its native range is to the north of its current introduced range. We then compared the SDM predictions to current observations of spread. Consistent with predictions, *T. clavata* appears to be spreading faster to the north than to the south. Lastly, we conducted surveys to investigate potential ecological impacts of *T. clavata* on the diversity of native orb weaving spiders. Importantly, *Trichonephila clavata* was the most common and abundant species observed in the survey, and was numerically dominant at half of the sites it was present in. Our models also suggest that there is lower native orb weaver species richness and diversity closer to where *T. clavata* was initially found and where it has been established the longest, though human population density complicates this finding. This early study is the first to forecast how widely this spider may spread in its introduced range and explore potential ecological impacts of *T. clavata*, and we call for continued investigation of this invasion’s effects.

## Introduction

Biological invasions are one of the most influential drivers of global change, with economic costs rivaling natural disasters (Turbelin et al. 2023). Across all biomes and systems, invasive flora and fauna can directly and indirectly affect native species, change population phenotypic and genotypic diversity, disrupt trophic networks, alter ecosystem function, drive species extinctions, and change the evolutionary trajectory of species (Brooks et al. 2004; Kenis et al. 2009; Ricciardi et al. 2013; Blackburn et al. 2019; Vitousek 1990). These complex and often significant impacts necessitate efforts towards early detection, evaluation, and if necessary, management of non-native species. Established monitoring programs exist for some non-native pest groups (e.g., Rabaglia et al. 2019) but there is no standardized method of population monitoring for most invasive species.

Monitoring the spread and impacts of non-native species is essential, but often difficult, because their effects on ecosystems may be weak at low initial population densities (Spear et al. 2021). Furthermore, many invasive species may be inconspicuous at low densities, leading to a significant time lag between species establishment and detection. By the time these species are detected, eradication and management options may be highly limited, though this can vary from one ecosystem to the next (Strayer 2020). When a conspicuous non-native species is detected early after introduction, it can provide a unique opportunity to evaluate its impacts as it spreads across the landscape.

The Jorō spider, *Trichonephila clavata* (L. Koch, 1878, Araneae: Araneidae), is a large-bodied orbweaver native to Asia, likely introduced to northern Georgia, U.S., around 2010 (Hoebeke et al. 2015, Chuang et al. 2023). The exact method of introduction is unknown but their arrival as stowaway hitchhikers on international shipping cargo is the most likely scenario (Hoebeke et al. 2015). Joro spiders have now spread into at least four states in the southeastern U.S., with breeding populations likely occurring in several additional states (Chuang et al. 2023, www.inaturalist.org). The rate at which these non-native arthropods are spreading appears to be increasing exponentially (Chuang et al. 2023), following a lag time typical of invasive species (Sakai et al. 2001), with their presence in at least ∼120,000 km^2^ as of summer 2022 (Chuang et al. 2023).

There is widespread interest and speculation on how far *T. clavata* may spread across North America. Given how recently this species was introduced to the continent, its native range may provide important clues to its future potential geographic distribution. *Trichonephila clavata* inhabits more northern latitudes in its native range than in its introduced range, which supports the potential for further northward spread. Despite their limitations (Liu et al. 2020), species distribution models (SDMs) can be a useful tool for providing future range predictions for invasive species and optimizing control and management strategies (Jiménez-Valverde et al. 2011; Kramer et al. 2017; Tingley et al. 2017; Giljohann et al. 2011; Tulloch et al. 2015). These models utilize species occurrence data, which can be derived from public monitoring efforts.

The sudden ubiquity of *T. clavata* in the southeastern U.S. coupled with their large size, bright coloration, and propensity to dwell around homes and other structures has brought these non-native spiders to the forefront of public awareness. In particular, the crowdsourced biodiversity observation site, iNaturalist (www.inaturalist.org), has been used as a platform for community and professional scientists, naturalists, and laypeople to upload photographs, verify identifications, and map distributions of several non-native pests worldwide (e.g., Werenkraut et al. 2020, Della Rocca and Milanesi 2022, Martel et al. 2022). There are over 6,700 “research grade” (i.e., including a photograph and multiple user verifications of species identity) observations of *T. clavata* globally as of May 2023, nearly half of which are from the non-native range. These observations first allowed us to make preliminary estimates of its range boundaries (Chuang et al. 2023), and here we further use them to construct a SDM to determine future range predictions and assess how it compares to *T. clavata*’s current range expansion rate.

*Trichonephila clavata*’s large size may facilitate detection, which can aid monitoring its ecological impacts as it spreads, despite the relative recency of invasion. Although invasive spiders have historically received little attention, case studies have demonstrated the ability of certain invasive spider species to compete with, displace, and prey on native spiders (Bednarski et al. 2010; Houser et al. 2014; Nyffeler et al. 1986; Coticchio et al. 2023) which can shift spider community composition (Jakob et al. 2011). *Trichonephila clavata* has been observed in locally high abundances, which may limit available web spaces for native orb weavers. At longer-established sites in Georgia, *T. clavata* has seemingly become the most common web-building spider (Davis and Frick 2022; DRC, DRN, JFD, MIS, pers. obs.). This raises questions of whether these densities come at the cost of native orb weaver abundance and diversity, especially with the native congener *Trichonephila clavipes* (Linnaeus, 1767).

We address standing questions on the nascent invasion of *T. clavata* in this study by: 1) building SDMs based on environmental features of their native range to predict the potential future spread of *T. clavata* in North America; 2) using SDM predictions to inform current range expansion dynamics; 3) using transect survey data to determine the effects of *T. clavata* presence on the abundance and diversity of orb weaving spiders in a portion of the southeastern U.S.

## Methods

### Objective 1

Building SDMs based on environmental features of *T. clavata*‘s native range to predict the potential future spread in North America.

#### Data and Data Cleaning

We downloaded all available research-grade records of *T. clavata* observations from both iNaturalist and GBIF as of November 10, 2022. We split these data into two regions: Asia and North America. We also downloaded the bioclimate dataset (19 variables) and wind speed from the WorldClim database at a resolution of 2.5 arc-minutes (∼4 km) (Fick and Hijmans 2017) and refer to these as bioclimate predictors hereafter. We then calculated a maximum wind speed variable based on the wind speed variable provided by WorldClim, following Segura-Hernández et al. (2023). We used QGIS to trim the original bioclimate predictor files to an Asian (52.083, -7.042: 146.833, 52.375) and North American (-125.042, 23.833: -64.417, 61.792) extent, respectively. We filtered *T. clavata* observations to only allow one observation per 4 km^2^ grid (spatial thinning) to control for biased sampling (Kiedrzyński et al., 2017; Steen et al., 2021) inherent in community-science data. We also randomly selected 50,000 background points (Valavi et al. 2022) for both Asia and North America and, along with our reduced observations, created two datasets that included the bioclimate predictors at each location.

#### Comparison of Asian and North American bioclimate predictors

We compared the climate (20 bioclimate predictors) of *T. clavata*’s native range to its introduced range using box plots, Mann-Whitney U tests, and the associated rank biserial correlation measure of effect size.

#### Bioclimate predictor reduction

We first tested the predictive value of each bioclimate predictor using bivariate logistic regression models. We then ranked each predictor based on the odds ratios derived from the models. Correlation analysis was then used to determine the degree to which each bioclimatic predictor was correlated with all other predictors. Highly correlated pairs of predictors (Pearson’s r > 0.7) were then identified. An iterative process was then used to sequentially remove the lowest performing predictors involved in any highly correlated pair until none were left. This reduced our bioclimate predictor dataset from 20 (19 bioclimate variables and wind speed) to five: minimum temperature of the coldest month, mean temperature of the wettest quarter, precipitation seasonality (coefficient of variation), precipitation of warmest quarter, and wind speed.

#### Creation of Species Distribution Models

After centering and scaling each predictor, we utilized four methods to create species distribution models trained on the Asian data. We used two high-performing machine learning approaches (a down-sampled random forest and maximum entropy; Liaw and Wiener 2002, Hijmans et al. 2011, and two statistical approaches (GLM and GAM; Wood 2017). We chose these methods following the suggestions of Zhu et al. (2020) and Valavi et al. (2022). Model creation was done using the *MGCV* (version 1.8-42, GAM and GLM), *dismo* (version 1.3-14, MaxEnt), and *randomForest* (version 4.7-1.1) packages in R. Data, R scripts, and other associated files can be found at 10.5281/zenodo.8091991.

To assess model overfitting, we performed 10-fold cross-validation on each of the four model types. Because conventional random cross-validation methods can underestimate prediction error for spatial data, we used the blockCV package (version 3.1-3) to assign folds using a spatial blocking technique (Valavi et al., 2019). We then used all available data to construct the final models. We used the predictions from these four methods and created an averaged model using QGIS (version 3.26.3). We report all five SDM models in the results to show their variation.

### Objective 2

Using SDM predictions to inform current range expansion dynamics.

#### Distance (Method 1)

We calculated the distance of every North American *T. clavata* record from the centroid of invasion. We calculated the invasion centroid by averaging the nine longitude and latitude coordinates from Hoebeke et al. (2015). We classified every observation as being NE, SE, NW, or SW from the centroid of invasion. To approximate the leading edge of *T. clavata* range expansion, we selected the 3 farthest observations in each direction from the centroid each year from 2018 to 2022. We calculated the mean value of the 3 selected observations and used this as a measure of the distance of range expansion (km) in each direction. [There was only 1 observation in 2018 in the SE direction.]

#### Area (Method 2)

We calculated the area (sq km) in North America occupied by *T. clavata* in each year from 2018 to 2022 using kernel density estimation (KDE) via the *amt* R package (version 0.2.1.0, Signer et al. 2019). To control for spatial bias in iNaturalist observations (i.e., observations are more heavily focused near large cities), we spatially thinned (as described above) *T. clavata* observations within each year. We performed KDE at 0.99 isopleth level at the reference bandwidth (h = (0.5*(sd long + sd lat)) * n^-⅙^). We then overlaid each year’s range on to four quadrants (NE of centroid, SE of centroid, NW of centroid, and SW of centroid) and calculated the area (sq km) of *T. clavata*‘s range in each of these four directions from 2018 to 2022.

### Objective 3

Determining the effects of *T. clavata*‘s presence on the abundance and diversity of orb weaving spiders with transect surveys

#### Transect data collection

Between August 29 and November 8, 2022 we surveyed 103 locations across northern Georgia, U.S. (Figure 1). We started a transect either at a location where *T. clavata* had never been observed previously, or an area where *T. clavata* had been observed for at least three consecutive years. All researchers followed the same survey protocol. We used a method similar to a Pollard walk as visual census techniques have been shown to be effective for counting spiders with conspicuous webs (Lubin 1978). We first, located and photographed an orb weaving spider, then we started a 10-minute timer, during which we moved continuously at a slow walking pace recording the presence and abundance of orb weaving spiders, pausing only to photograph each spider to confirm their initial identification. We recorded the date, time, temperature, windspeed, and amount of precipitation within the last 24 h at each survey location.

**Figure 1:**
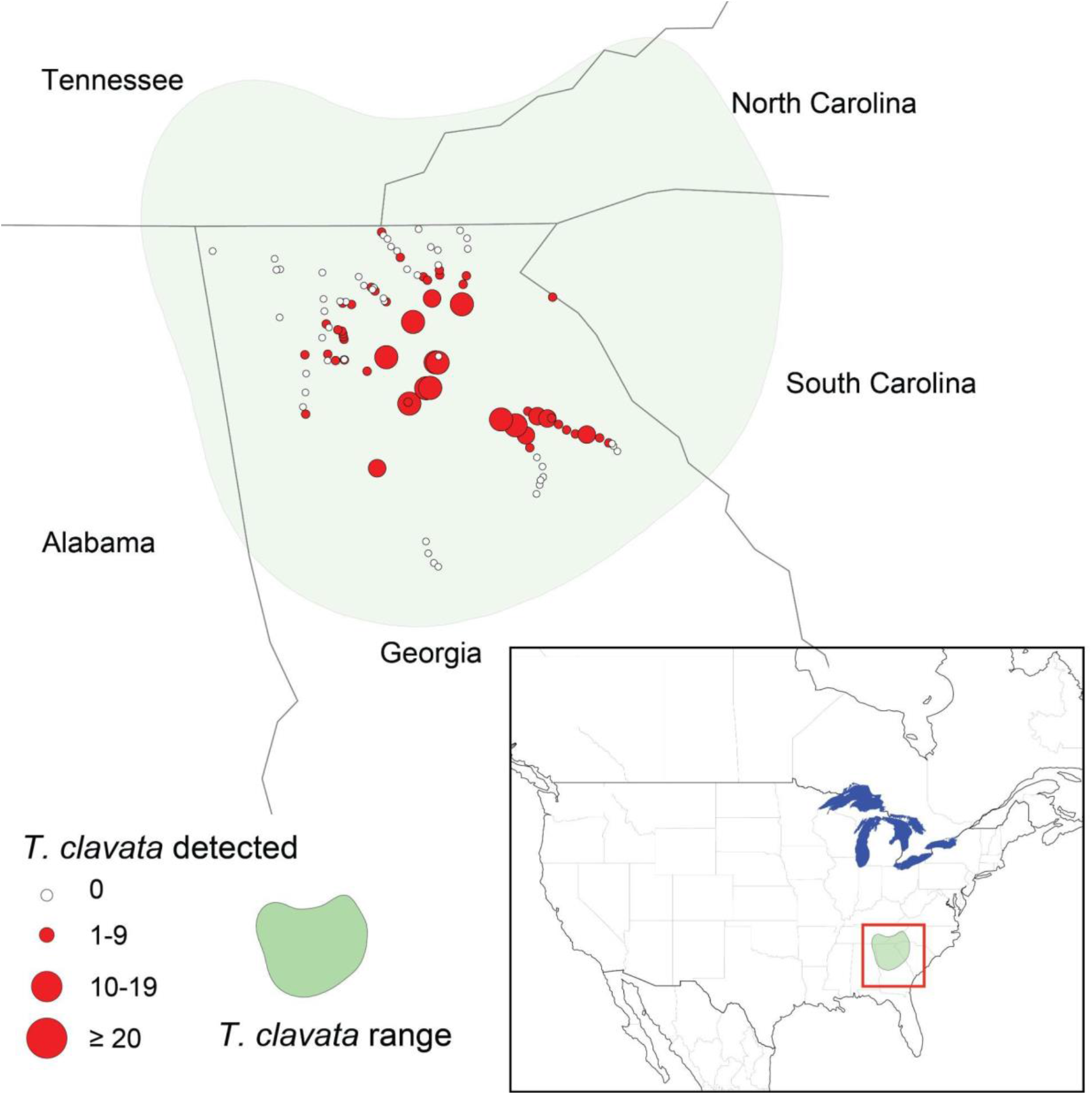
Map of survey transect locations where *T. clavata* and native orb weaver abundance was recorded. Abundance of *T. clavata* is indicated by the size and color of the circle. Range of *T. clavata* (using iNaturalist data through December 2022) is indicated by the shaded region. Inset map shows location of *T. clavata* range and survey transects in relation to North America.

#### Variable calculation

Following the recommendations of Mouillot and Lepretre (1999) and Morris et al. (2014) we used several measures of species diversity, including species richness, Shannon–Weaver’s index, and Simpson’s diversity index as our outcome (dependent) variables. We used the vegan package (version 2.6-4) in R (Oksanen et al. 2022) to calculate Shannon’s and Simpson’s diversity indexes. We calculated each measure of diversity across all locations independent of the abundance of *T. clavata* at that location. We did this because we were interested in how the presence of *T. clavata* affected the diversity of native orb weavers and whether the inclusion of *T. clavata* altered species richness and/or species evenness, especially when the density of *T. clavata* was high.

For our predictor (independent) variable, we estimated two measures of the historic presence of *T. clavata* across our survey area: 1) distance from the invasion centroid to each surveyed location (described above) and 2) an estimate of how long *T. clavata* has been present at a surveyed location. We calculated the number of years *T. clavata* had been in an area by creating a 10 km by 10 km grid, noting the date of the first observation within each cell, then calculating the number of years from that time until November 2022. We assigned a value of 1 year to cells with a first observation in 2022, to account for the presence of *T. clavata* during the summer of 2022 prior to field surveys. We assigned values to survey locations according to the grid cell within which they are located. Lastly, because the introduction of *T. clavata* occurred around the greater Atlanta metropolitan area, we controlled for urbanization by obtaining the human population density of each surveyed location from the “Gridded Population of the World (GPW), v4” dataset (CIESIN 2018) at a resolution of 30 arc-seconds (1 km).

#### Data Analysis

We tested for spatial autocorrelation between survey locations using the spdep package (version 1.2-8) in R (Bivand et al. 2013, Bivand 2022). We constructed a spatial correlogram to visualize and explore our data. We also calculated Moran’s I for our three diversity indexes using two approaches. First, we ensured that all locations had at least one neighbor. Second, we applied a 10 km radius around each location, all locations that fell within that 10 km radius were considered neighbors. The spatial correlogram showed no clear spatial pattern and none of the Moran’s I results were significant. Following these results, we used both linear and generalized linear models to statistically test for the impact of *T. clavata* presence on the species richness and diversity of native orb weaving spiders.

To aid in the construction of biologically relevant statistical models, we constructed a directed acyclic graph (DAG) to represent our proposed causal relationship between the predictor and outcome variables and identify potential confounders (Supplemental Figure A). Although DAGs are very useful in model creation, they are of limited use for determining which interactions to include in the final models (Attia et al. 2022). Therefore, we created several versions of each model, some with only main effects and some including interactions. We used AIC/R^2^ values to determine which model to focus on for our final interpretations. We constructed regression models for each combination of *T. clavata* predictor variables (distance from the invasion centroid and years occupying a location) and outcome variables (species richness, Shannon’s index, and Simpson’s index). We used multivariate linear regression when Shannon’s and Simpson’s diversity indexes were the outcome variable respectively and multivariate Poisson regression when species richness was the outcome variable. Model equations followed the following format: outcome variable ∼ Temp (C°) + Windspeed (kph) + Rain in the last 24hr (cm) + Total days of the year (out of 365) + Population density*predictor variable. To avoid the “table 2” fallacy, we only report the model coefficients for our predictor variables and not for any of the potentially confounding variables identified by our DAG (Westreich and Greenland 2013).

## Results

### Comparison of Asian and North American Bioclimate Predictors

All 20 predictor variables differed significantly between Asian and North American windows (Figure 2, Supplemental Table A), as most climate predictors showed a large effect (*r*_rb_ > 0.5) for differences by region. However, mean temperature of the warmest quarter, temperature seasonality, annual mean temperature, and wind speed showed only moderate effects (*r*_rb_ > 0.3), and minimum temperature of coldest month, annual precipitation, and annual temperature range showed small effects (Figure 2, Supplemental Table A).

**Figure 2:**
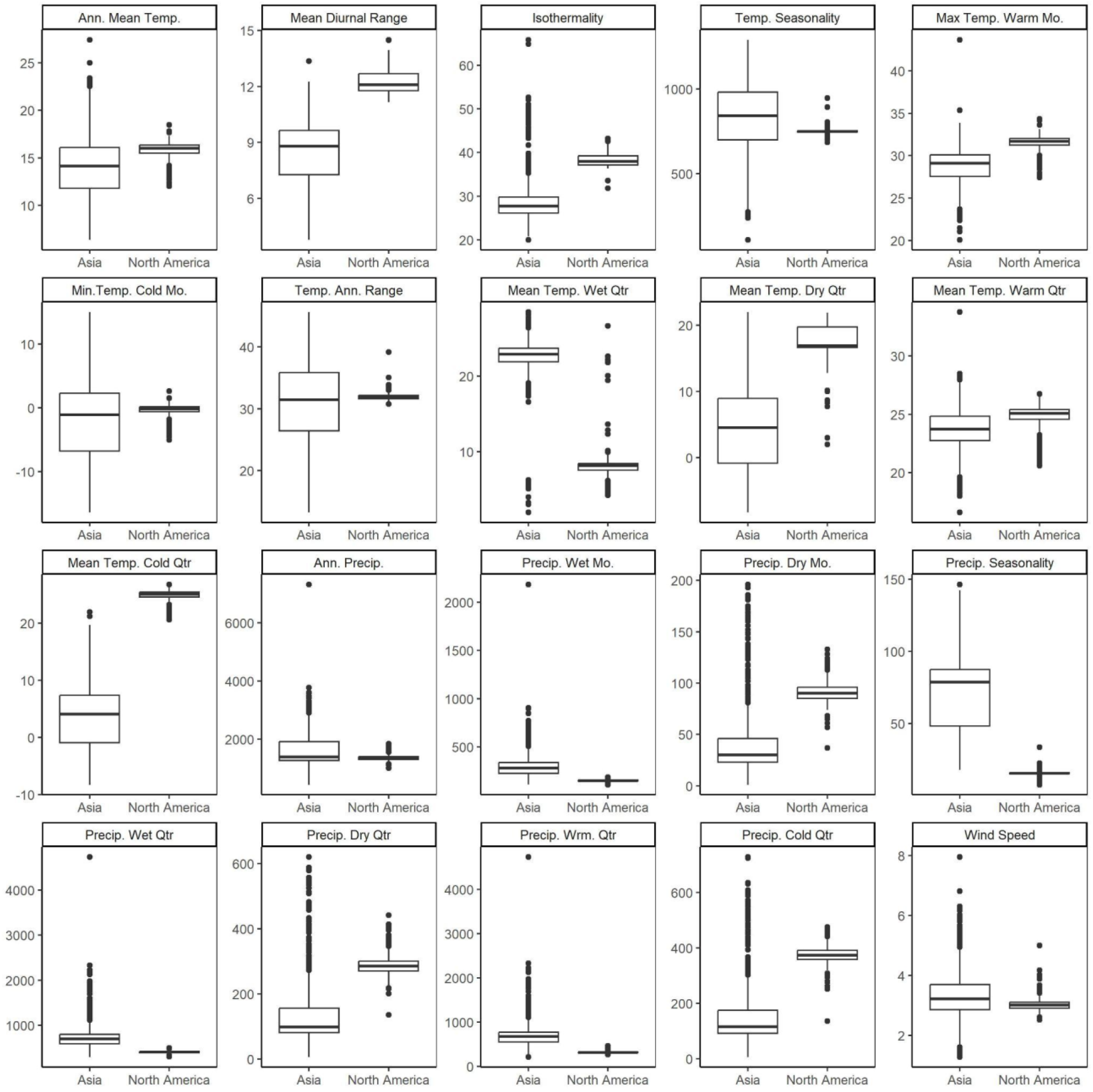
A comparison of the values of the 19 standard bioclimatic variables and wind speed for each of the known ranges of *Trichonephila clavata* in both Asia and North America. Units for temperature related variables are in °C except for *Temp. Seasonality* which is in °C·100. The unit for precipitation variables is kg·m^-2^. Wind speed is in m·s^-1^. Spider observations from iNaturalist (https://www.inaturalist.org/) and bioclimatic data from WorldClim (https://worldclim.org/).

These results show that total annual precipitation between known *T. clavata* locations in Asia and North America are similar, but the pattern of when this precipitation falls varies. Asian T*. clavata* experiences a greater seasonal fluctuation in rainfall than those in North America (i.e., Asian spiders experience more distinct rainy and dry seasons). Likewise, temperature seasonality and the annual temperature range are similar between locations where *T. clavata* is found in Asia and North America; however, how the temperature fluctuates throughout the year varies. North American spiders experience greater daily fluctuations in temperature (max temperature in the warmest month and mean diurnal range) and are warmer in the coldest quarter and driest quarters, while Asian spiders experience warmer conditions in the wettest quarter.

### Species Distribution Models

*Trichonephila clavata* has become established in the southeastern U.S., and our models show that this region differs climatically from *T. clavata*’s native range. Based on SDM models that performed well in predicting the Asian distribution, our SDMs suggest that the areas of North America that are most similar to its native habitat are predominantly further north than the areas *T. clavata* currently occupies. Ten-fold cross-validation results ranged from Mean AUC_ROC_ = 0.77 – 0.92, with the GLM performing the worst and all other models performing equally. The performance of each model is reported in Figure 3. Our application of these models to North America suggests that the Great Lakes region of the U.S. and Canada extending throughout the midwestern and northeastern U.S., and into eastern Canada, are most similar to *T. clavata*’s native range in Asia (Figure 3). Our models also show that areas of the American northwest (both the U.S.and Canada, all four models) and the mountainous region of northwestern Mexico (GLM, MaxEnt, and RandomForest, but not the GAM) may also have climates similar to that in *T. clavata*’s native range. Our ensemble (averaged) model suggests that *T. clavata* may be capable of surviving in a substantial portion of North America, from its current distribution west to the Great Plains and north into Canada, and in areas of the Pacific Northwest in the U.S. and Canada.

**Figure 3:**
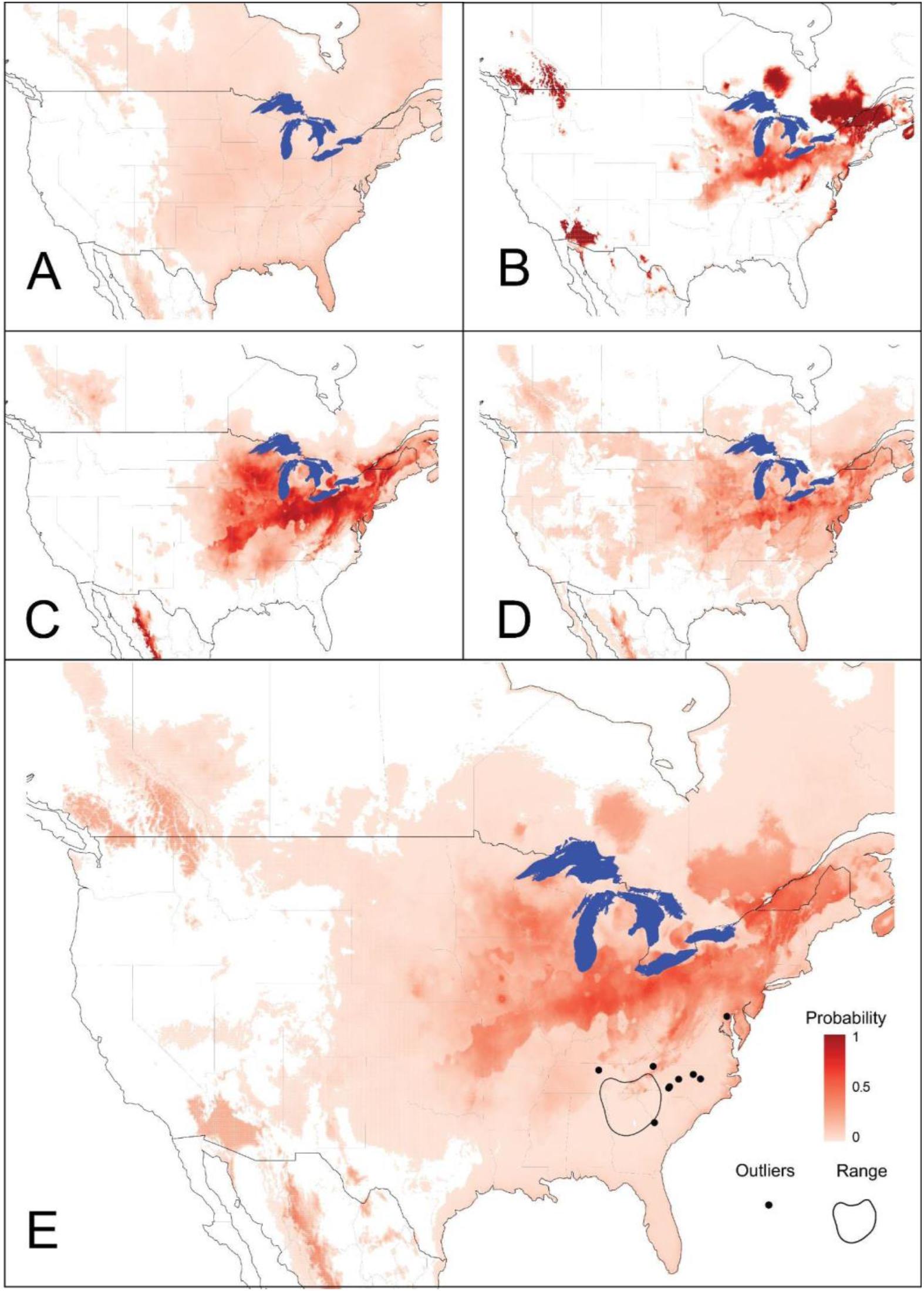
Species distribution model (SDM) predictions of *Trichonephila clavata*’s suitable range in North America based on its native Asian distribution: **A)** General Linear Model: cross-validation Mean AUC_ROC_ = 0.77 and AUC_ROC_ of model shown = 0.89, **B)** General Additive Model: cross-validation Mean AUC_ROC_ = 0.92 and AUC_ROC_ of model shown = 0.99, **C)** Maximum Entropy: Mean AUC_ROC_ = 0.92 and AUC_ROC_ of model shown = 0.96, **D)** Random Forest: Mean AUC_ROC_ = 0.92 and AUC_ROC_ of model shown = 0.99, and **E)** Averaged prediction of all four models, calculated as an equally weighted average across all four primary models, including *T. clavata’s* introduced range as of December 2022.

We compared SDM predictions with current *T. clavata* range expansion dynamics as captured by iNaturalist data. Whether quantified by area or distance, the data indicate that *T. clavata* is spreading faster to the north than to the south (Figure 4). *Trichonephila clavata* has spread 436 and 245 km to the NE and NW, respectively, compared to 113 and 131 km to the SE and SW.

**Figure 4:**
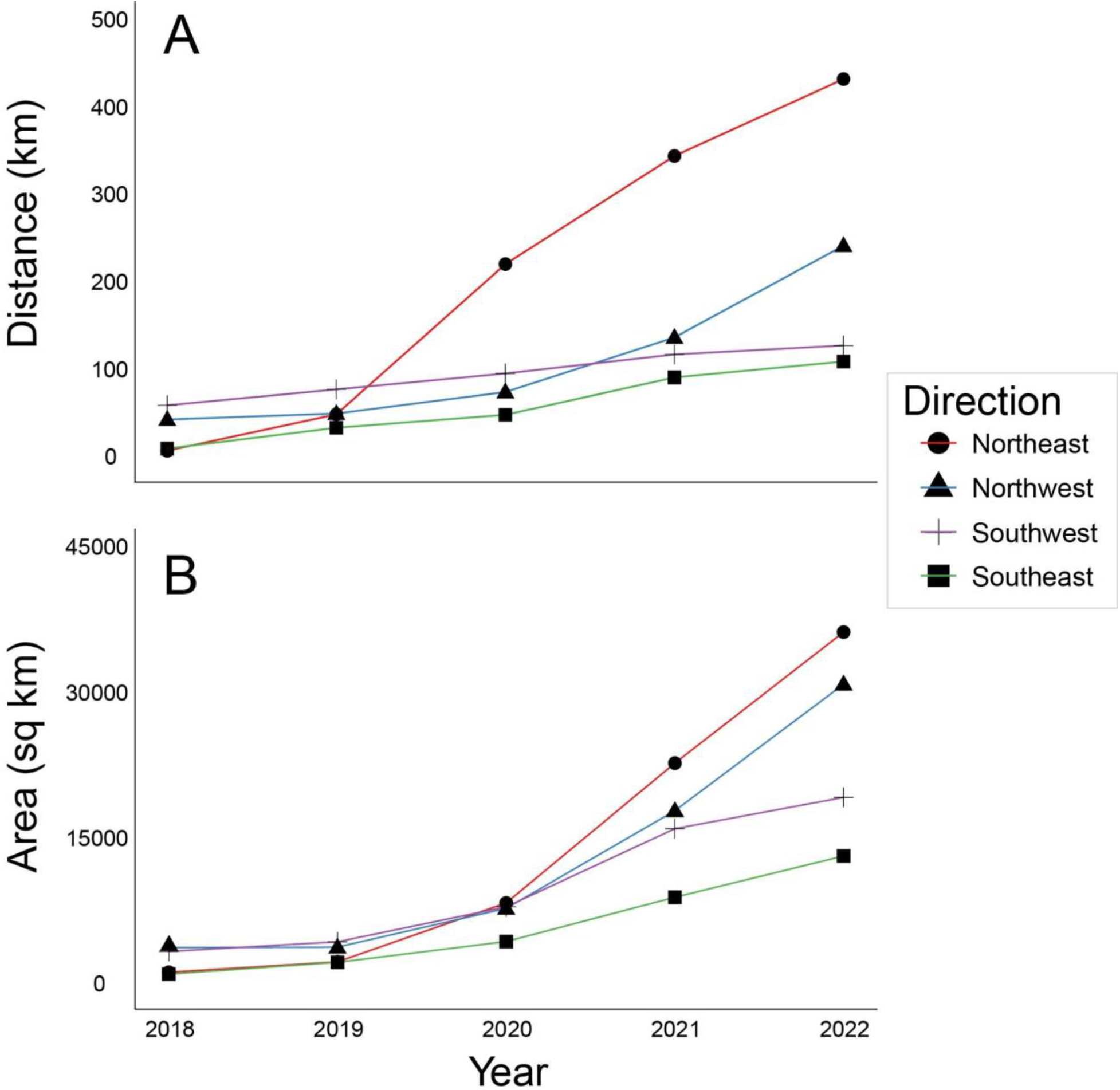
Rate of range expansion by *Trichonephila clavata* from 2018 through 2022, measured by **(A)** distance (km) of range edge from point of introduction and **(B)** area (sq km).

### Descriptive Results of the Survey

We found 19 different orb weaving spider species on our transect surveys, though spider abundance was numerically dominated by *T. clavata* and the native *Micrathena mitrata* (Figure 5; Supplemental Table B; Supplemental Figure B). *Trichonephila clavata* (n = 543) was observed at the most locations (53 of 103, or 51.5%), was the most abundant spider observed on average (>5 individuals per location on average) and was the species with the most individuals found in total across all surveyed locations. At half of the sites where it was present, *T. clavata* was the most numerically dominant species. *Micrathena mitrata* (n = 319) was the next most abundant, found in 49 of 103 locations (47.6%) with an average of >3 individuals per site. Only one other species (*Neoscona crucifera*) was observed at >40% of surveyed locations (44 of 103 or 42.7%), and it, along with all other species, was less abundant (<2 individuals on average per site) overall. The total number of individuals observed for all species other than *T. clavata* and *M. mitrata* (n = 386) was less than *T. clavata* alone and only slightly larger than *M. mitrata*.

**Figure 5:**
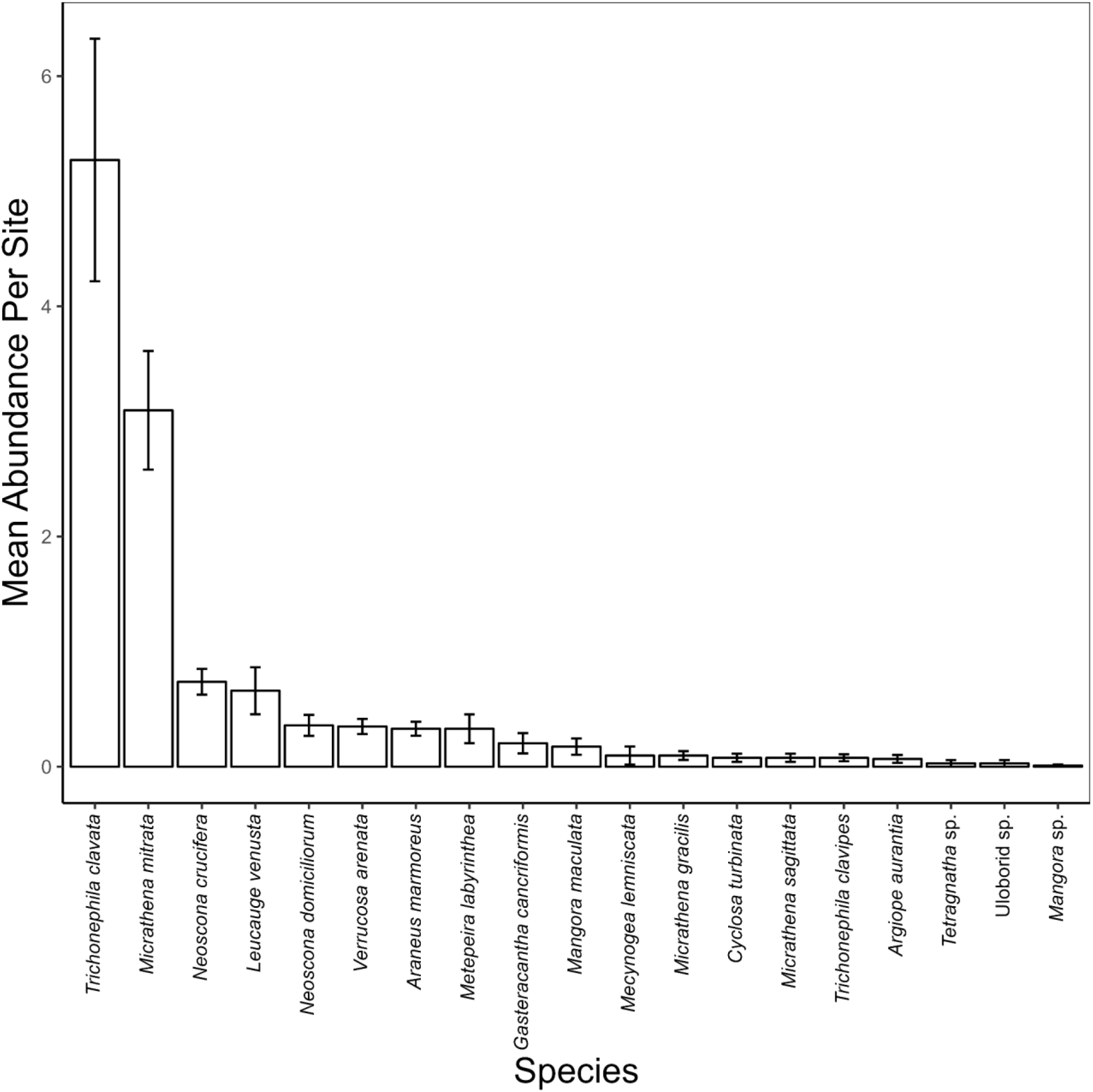
Mean abundance (+/- SE) of spider species observed across all surveyed locations.

### Impact of *T. clavata* on Native Orb Weaving Spider Species

We found evidence that the length of time *T. clavata* has been present at a location was typically a significant predictor of orb weaving spider diversity at that location (Table 1). We found consistent evidence that the models including the interaction between historical presence of *T. clavata* and human population density performed better than the other models as assessed by AIC scores and R^2^ values. Thus, we focus our attention on these models for the remainder of the results and discussion. In addition, we only present the graphical results for species richness and Shannon’s index as the results for Simpson’s index (see Supplemental Figure C) were consistent.

**Table 1:** Results of linear and general linear models testing if the presence of *Trichonephila clavata* affects the abundance and diversity of native orb-weaving species. This table reports the coefficients, SE, P-value, and AIC/R2 for all linear models: main effects not including human population density, main effects controlling for human population density, and the interaction between human population density and *T. clavata’s* historical presence.

| Outcome | Historic Predictor | Coefficient ± SE |  |  | P |  |  | AIC/R<sup>2</sup> |  |  |
| --- | --- | --- | --- | --- | --- | --- | --- | --- | --- | --- |
|  |  | Excluding Pop | Pop Density | Interaction | Excluding Pop | Pop Density | Interaction | Excluding Pop | Pop Density | Interaction |
| Species Richness | Distance from Centroid | 4.1E-6±2.2E-6 | 3.6E-6±2.4E-6 | 2.9E-8±1.3E-8 | 0.062 | 0.14 | **0.032** | 377 | 378.8 | **376** |
|  | Years at Location | -0.059±0.044 | -0.044±0.050 | -6.6E-4±2.8E-4 | 0.17 | 0.38 | **0.016** | 378.6 | 380.2 | **375.7** |
| Shannon's | Distance from Centroid | 6.1E-6±1.9E-6 | 4.4E-6±2.1E-6 | 2.1E-8±1.0E-8 | **0.0016** | **0.037** | **0.039** | 0.17 | 0.2 | **0.24** |
|  | Years at Location | -0.099±0.037 | -0.059±0.042 | -4.0E-4±1.8E-4 | **0.0094** | 0.16 | **0.033** | 0.15 | 0.18 | **0.22** |
| Simpson's | Distance from Centroid | 2.3E-6±1.1E-7 | 1.4E-6±9.2E-7 | 9.2E-9±4.5E-9 | **0.0076** | 0.13 | **0.043** | 0.1 | 0.14 | **0.17** |
|  | Years at Location | -0.035±0.016 | -0.016±0.018 | -1.7E-4±8.0E-5 | **0.033** | 0.4 | **0.034** | 0.08 | 0.12 | **0.16** |

Distance coefficients are in meters
Pearson's Corr Distance x Population: P = 3.3E-5, Pearson's = -0.40 (-0.55 – -0.22 95% CI)
Pearson's Corr Years at Location x Population: P = 1.2E-7, Pearson's = 0.49 (0.33–0.63 95% CI)

Our models suggested lower orb weaving spider diversity in areas where *T. clavata* had been present for a longer period of time and greater orb weaving spider diversity in areas further from the invasion centroid. However, the significant interactions in these models suggested that these effects were stronger in areas with larger human populations. Models where the human population was held constant at the mean or the mean + 1 SD for all survey sites predicted higher diversity and richness than models held constant at the mean human population - 1 SD as the distance from invasion centroid increased (see Figure 6A & C).

**Figure 6:**
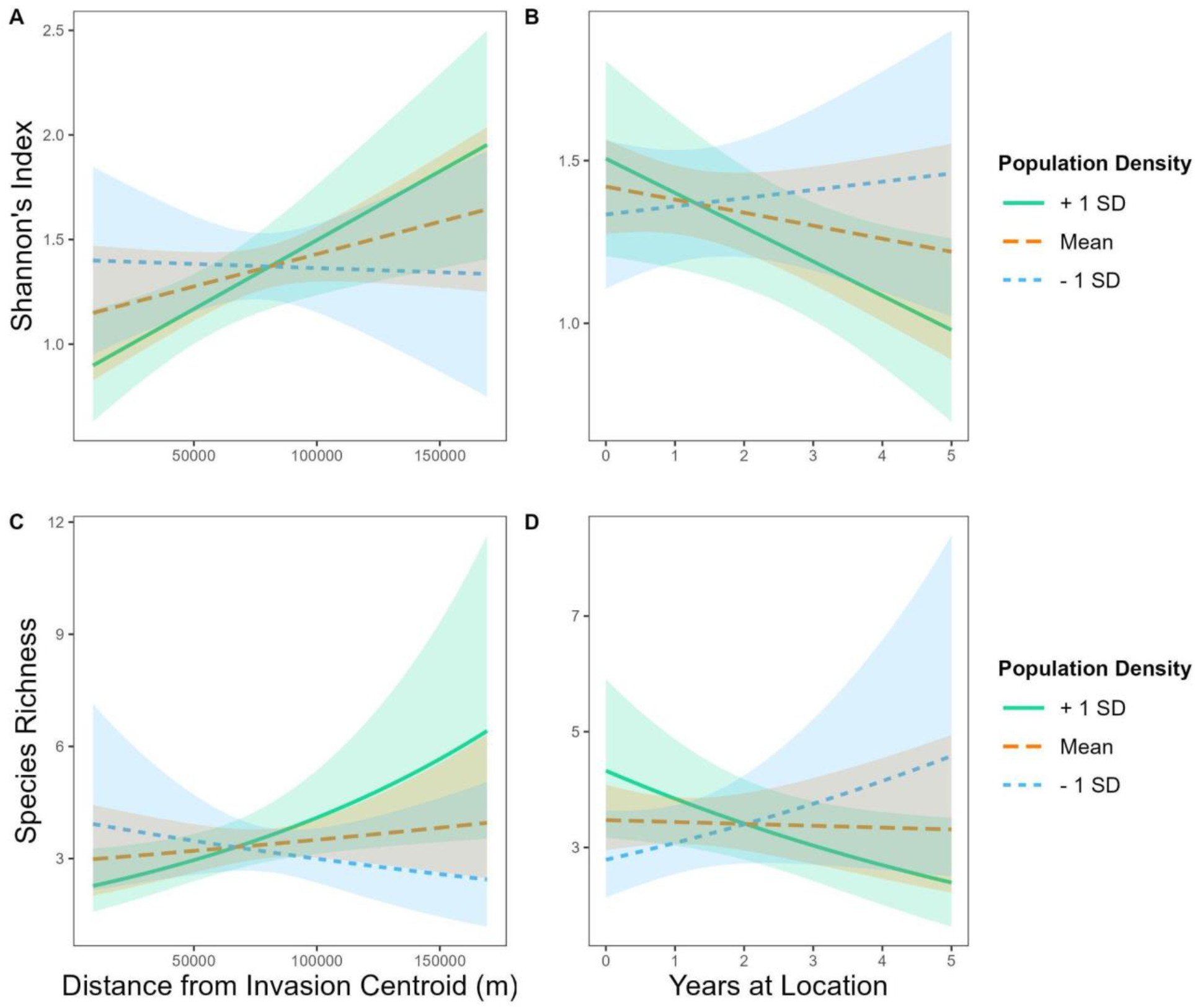
Interaction plots based on linear and generalized linear regression models of a location’s human population density and a measure of *Trichonephila clavata’*s presence (in space or time) on orb weaving spider diversity. Shading around trend lines represents 95% CI.

Specifically, holding human population density constant, for every 1 km increase in distance from the invasion centroid we expect a 2.9 x 10^-5^ increase in the number of species present and 2.1 x 10^-5^ increase in the Shannon’s diversity score. Thus, we expect a site with + 1 SD and located at the invasion centroid to have 2.76 species present and a Shannon’s score of 1.21, while a site 100 km away and all other variables held constant would see an increase in species richness (5.95) and Shannon’s diversity score (1.77). Likewise, models where the human population was held constant at the mean or the mean + 1 SD predicted lower diversity and richness than models held constant at the mean human population - 1 SD as the number of years *T. clavata* occupied a site increased (see Figure 6B & D). For example, at sites where human population density is held constant, with every 1-year increase in the length of time *T. clavata* have been present we expect a 6.6 x 10^-4^ decrease in the number of species and a 4.0 x 10^-4^ decrease in Shannon’s diversity score. Therefore, we expect site with + 1 SD population density and where *T. clavata* has never been present to have 3.78 species present and a Shannon’s score of 1.43, while a site that had *T. clavata* for 5 years but maintaining the same human population density would show a decrease in species richness to 1.56 and a Shannon’s diversity score of 0.90.

## Discussion

### Species Distribution Models

Overall, our results suggest that *T. clavata* may be capable of significant range expansion within North America. Our species distribution models, basing *T. clavata’s* optimal range on its native Asian distribution, found the strongest habitat similarity in regions south of the Great Lakes and all states in the northeastern USA. All four models also predicted portions of Canada, including provinces north of the Great Lakes, such as Quebec and Ontario, as well as more fragmented portions of Alberta and British Columbia as suitable matches. Lastly, the models suggested fragmented sections of Mexico, near Chihuahua, as more similar to climates *T. clavata* inhabits in Asia. These areas are all currently outside of *T. clavata*’s introduced range, though ongoing spread may eventually result in their presence in these distant regions.

Our GAM and MaxEnt models predict no, or almost no, climate similarity where *T. clavata* is apparently thriving in the southeastern USA, while our RandomForest and GLM models predicted these areas as having only low similarity. This may be due in part to the relative differences between the bioclimate predictors for Asia and North America. Not only did many of our predictor variables have a large difference between regions, but three (mean temperature of wettest quarter, precipitation seasonality, and precipitation of warmest quarter) of the five variables used in our SDMs had large differences, while the remaining two variables (maximum wind speed and precipitation of warmest quarter) had moderate and small differences, respectively. In their North American range, *T. clavata* experiences lower mean temperatures during the wettest quarter, lower precipitation seasonality, and lower precipitation during the warmest quarter.

All of this suggests that models trained on data from Asia may have limited application when used to predict the general suitability of North America for *T. clavata’s* expansion. However, despite this limitation, these models still offer insights into the possible range expansion of *T. clavata* throughout North America. First, similar to Luo et al. (2022), and following the recommendation of Hui (2023), we limit our interpretation of our model results to a comparison of climatic similarity between Asia and North America. Our results suggest that *T. clavata* may have experienced a shift in its realized niche, similar to what Luo et al. (2022) found with *Latrodectus hasselti* (which is native to Australia) in its introduced range of Japan, New Zealand, and Southeast Asia. Second, although all climatic variables differed significantly between Asia and North America, the differences between minimum temperature, temperature range, and annual precipitation were small or very small (see Supplemental Table A). This suggests that areas of North America identified as most similar to *T. clavata*’s native Asian range have temperatures and total precipitation that this species is already capable of surviving.

Biotic interactions may explain why *T. clavata’s* latitudinal distribution in Asia climatically differs so much from its current introduced range. Native North American antagonists (e.g., birds, predatory wasps, and parasitoids) may not be regulating *T. clavata* populations as they do in Asia (i.e., the enemy release hypothesis; Roy et al. 2011, but see Powell and Taylor 2017). Competition should also be considered, but it is unknown whether smaller orb weavers are significant competitors, as *T. clavata* may capture larger prey that is unavailable to smaller spiders.

However, competition with other large-bodied orb weavers with correspondingly large webs is likely, especially between congeners that may share ecologically similar niches. *Trichonephila clavata*’s Asian range comprises Japan, the Korean peninsula, China, Taiwan, Myanmar, as well as the Himalayan regions extending to India (Figure 7). While *T. clavata’s* range overlaps with the more geographically common *Nephila pilipes* (Fabricius, 1793), *T. clavata* rarely occurs with *Trichonephila antipodiana* (Walkenaer, 1841) and *Nephila kuhli* (Doleschall, 1859). In fact, *T. antipodiana* and *N. kuhli* are only present where *T. clavata* is absent or rarely found in southern Myanmar, Thailand, Laos, and Cambodia (Figure 7); this pattern is reflected in iNaturalist occurrence data and on GBIF records (www.gbif.org).

**Figure 7:**
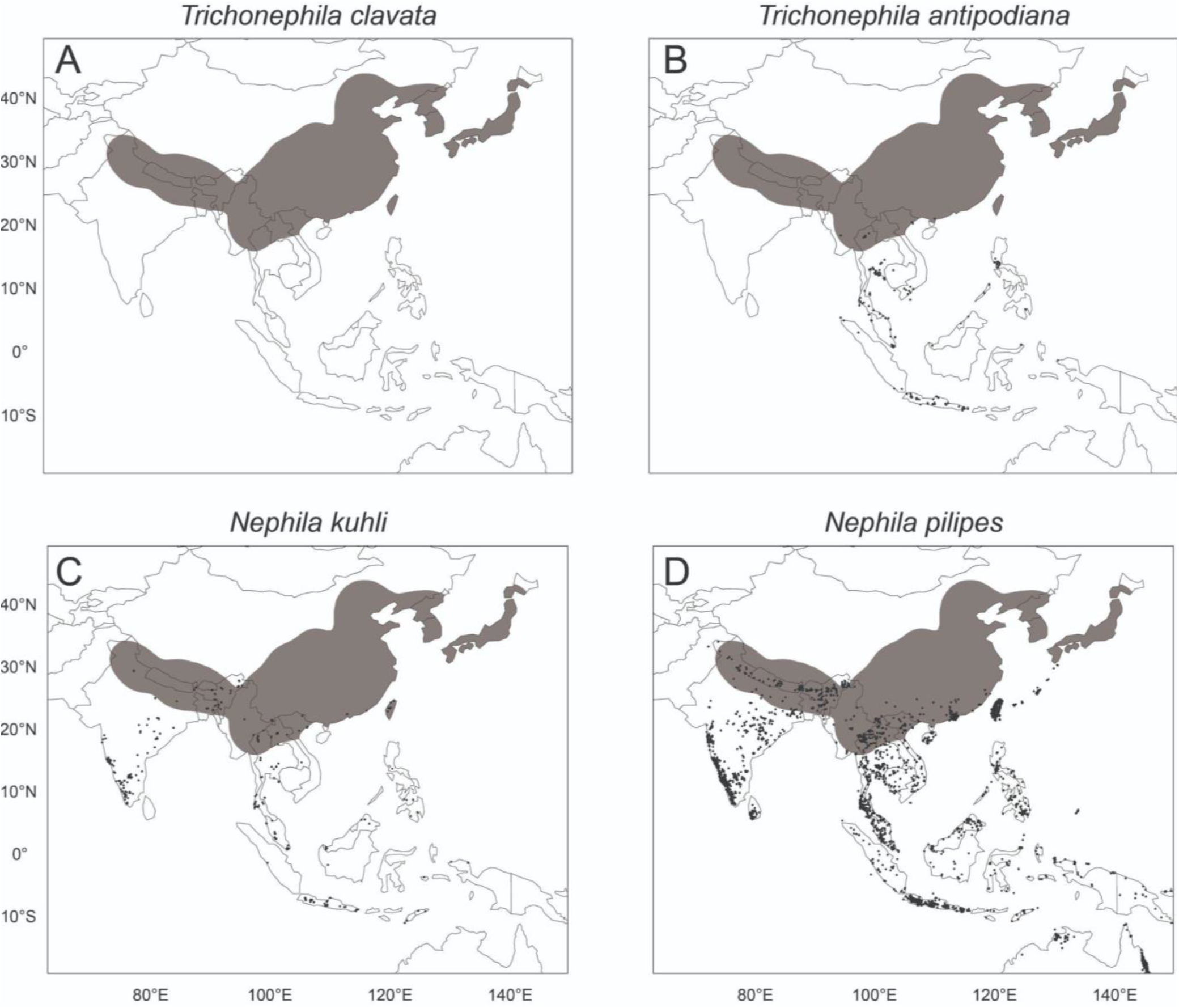
Distribution of A) *Trichonephila clavata* in its native range in comparison with occurrences of three related species, B) *Trichonephila antipodiana*, C) *Nephila kuhli*, and D) *Nephila pilipes* based on iNaturalist and GBIF records (www.inaturalist.org; www.gbif.org) as of May 2023. Gray shading represents *T. clavata*’s range in all panels. Darker points represent individual reports of other species.

This striking lack of overlap between *T. clavata* and these two species raises the possibility that competitive exclusion might restrict *T. clavata*’s realized niche. If so, *T. clavata* may have higher environmental flexibility than is suggested from its native range, which would help explain the discrepancy between its current establishment in the southeastern U.S. and the range predicted by our SDM models. Without any comparable large-bodied orb weavers to the north of *T. clavata*’s introduction area, range expansion may go largely unchecked by competition as an empty niche is filled. Interestingly, southern expansion may be slowed or limited by the presence of *T. clavipes*. These species currently overlap at the edges of their distributions, so detailed study of their interactions is needed.

It is also possible that *T. clavata’s* known Asian range reflects sampling bias, with fewer observations made in southeastern Asia. Sampling bias is an inherent risk when using data from crowdsourced websites like iNaturalist (Di Cecco et al. 2021). The relative absence of *T. clavata* observations in Myanmar and Vietnam as well as the rest of continental southeastern Asia raises questions of whether differences in the number of iNaturalist users and observations across countries might bias our models. However, the extensive geographic coverage of *T. clavata*’s relatives reported in these regions, especially *N. pilipes* (over 14,000 research grade observations on iNaturalist alone), suggests the relative absence of *T. clavata* in southeastern Asia in our datasets likely represents a real pattern rather than a bias attributable to a dearth of users or observations.

Our models suggest that climates in the northeastern USA and southern Canada are more suitable matches for *T. clavata* than their current introduced range, so cooler northern climates will likely not impede continued range expansion. In spiders, natal dispersal is typically achieved by ballooning, where newly emerged spiderlings release silk and travel as aeronauts in air currents (Foelix 1996). *Trichonephila* and *Nephila* spiders are no exception, as they are known for their long-distance ballooning abilities (Jung et al. 2006; Kuntner and Agnarsson 2011; Lee et al. 2015). This, paired with climatic suitability across eastern North America, may explain the observed range expansion dynamics. Our SDM predictions, coupled with *T. clavata*’s substantial powers of dispersal, limited competition with comparably sized spiders, and release from enemies in its native range, suggest continued expansion to the Northeast as an unoccupied niche is filled. Disentangling the mechanisms facilitating this range expansion is an area primed for future research.

### Impact of *T. clavata* on native orb weaving spiders

Our study reports the first known ecological impacts of *T. clavata* in its introduced range, specifically on native orb weaving spiders. We found reduced diversity of native orbweavers in areas with prolonged presence of *T. clavata* and higher human populations; however, since these models were based on data with few *T. clavata* observations in areas with very low human population density, this conclusion must remain tentative. Future studies should experimentally test for ecologically meaningful relationships between *T. clavata* and native orb weavers to thoroughly explore the impacts of the *T. clavata* invasion to supplement our first year of correlative observational data.

Another explanation is that *T. clavata* may be more tolerant of urban environments with greater human activity than some native species. Several spiders are more commonly found in areas with a greater human impact (e.g., Fraser and Frankie 1986, Moorhead and Philpott 2013, Damptey et al. 2022), and some species actively thrive in these areas due to increased reproductive capacity (Lowe et al. 2014) and enhanced prey capture (Gomes 2020). Many of the known North American non-native spiders are synanthropic (e.g., *Pholcus phalangioides* and *P. manueli*: Campbell et al. 2020; *Cyrtophora citricola*: Chuang and Riechert, 2021; *Latrodectus geometricus*: Vetter et al. 2016; *Oecobius navus*: Voss et al. 2007), likely due to selective pressures associated with their common introduction pathway as shipping cargo stowaways (Hanggi and Straub 2016; Nentwig 2015). While *T. clavata* shares this proposed introduction pathway (Hoebeke et al. 2015), it is unlike the other species because it is present in both urban and natural environments in its introduced range, which increases the odds of impacts on native communities. Ultimately, more information is needed to elucidate the interactions between non-native and native arthropods and urban areas (McIntyre 2000; Gardiner et al. 2013) as well as the potential impacts of this invasive species as it continues to spread.

Although it has only been documented in North America for less than fifteen years, *T. clavata* has already become the most locally common and abundant orb weaving spider species in our surveyed area. The rapid spread and numerical dominance classify *T. clavata* as an invasive species (*sensu* Richardson 2011). Invasive spiders can negatively impact native arthropod communities in many ways. For example, the presence of invasive *Linyphia triangularis* led to reduced densities of native spiders (Jakob et al. 2011), increased web abandonment and web building of the native *Frontinella communis*, and web takeovers of *F. communis* in Maine, U.S. (Bednarski et al. 2010). Candy-striped spiders (*Enoplognatha ovata* and *E. latimana*) are known to disproportionately prey on pollinators, even actively hunting prey during pre-dawn periods when insects are typically lethargic (Scott and McCann 2023). We have observed *T. clavata* feeding on prey in several different arthropod groups, including cockroaches, beetles, wasps, bees, butterflies, dragonflies, grasshoppers, and other spiders (DRC, JFD, MIS, pers. obs.). The high metabolic rate and quick maturation time of *T. clavata* (Davis and Frick 2022) suggests a substantial amount of prey is taken, and thus not available to native orb weavers. Detailed studies of interactions between *T. clavata* and native orb weavers and prey are needed to better understand community responses to this invasion.

## Conclusion

There is an abundance of suitable habitat for *T. clavata* throughout eastern North America and in some areas in the western part of the continent. With the large population expansion rate seen so far, coupled with both natural and anthropogenic dispersal mechanisms, it seems highly likely that *T. clavata* will continue to spread into these areas. *Trichonephila clavata* is often numerically dominant among orb weavers where it occurs, both more common and abundant than native species across this survey. Future surveys will continue exploring the nature of *T. clavata*’s relationships with native spider communities. In the meantime, we continue to advise adopting a measured approach surrounding the science, management, and reporting on this invasive spider.

## Supporting information

All Supplemental Files

## Data Availability

The data that support the findings of this study are openly available in Zenodo at DOI: https://doi.org/10.5281/zenodo.8091991.

