## Supplementary material for "Veni, vidi, vici? Future spread and ecological impacts of a rapidly-expanding invasive predator population": All Supplemental Files

Supplemental Figure A: Directed Acyclic Graph of the proposed causal relationship between predictor variables (population density, *Trichonephila clavata* presence, and their interaction) on the outcome variable (orb-weaver diversity). The outcome variable is indicated by the blue oval with an “I”, the predictor variables are indicated by the green ovals with “play” symbols, and the confounding variables are depicted as white ovals. The oval labeled “current abundance” is a mediator and therefore not included in the regression models.

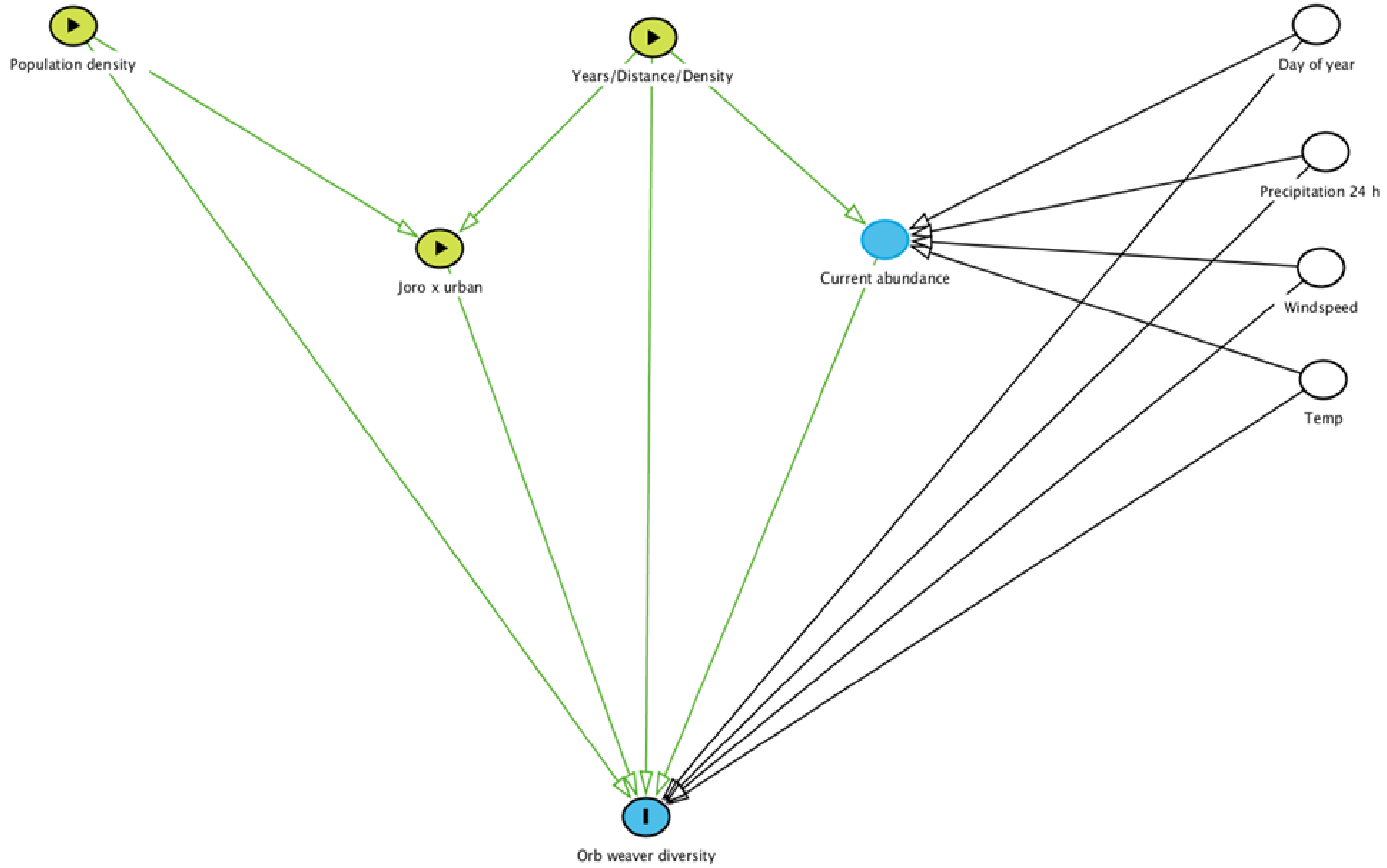

Supplemental Table A: Results of Mann-Whitney U test comparing bioclimate variables between Asia and North America.

| Variable | *W* | *p* | $r_{rb}$ | Effect size classification |
|---|---|---|---|---|
| Annual Mean Temperature | 1712432.5 | <0.0001 | -0.359 | Moderate |
| Mean Diurnal Range | 40296.5 | <0.0001 | -0.848 | Large |
| Isothermality | 443302 | <0.0001 | -0.731 | Large |
| Temperature Seasonality | 4174458 | <0.0001 | 0.36 | Moderate |
| Max Temperature of Warmest Month | 368615.5 | <0.0001 | -0.752 | Large |
| Precipitation of Warmest Quarter | 2536077 | <0.0001 | -0.119 | Small |
| Temperature Annual Range | 2708356 | <0.0001 | -0.0684 | Very Small |
| Mean Temperature of Wettest Quarter | 5850673 | <0.0001 | 0.851 | Large |
| Mean Temperature of Driest Quarter | 251011.5 | <0.0001 | -0.787 | Large |
| Mean Temperature of Warmest Quarter | 1480008.5 | <0.0001 | -0.428 | Moderate |
| Mean Temperature of Coldest Quarter | 27 | <0.0001 | -0.86 | Large |
| Annual Precipitation | 3346232 | <0.0001 | 0.118 | Small |
| Precipitation of Wettest Month | 5830043 | <0.0001 | 0.844 | Large |
| Precipitation of Driest Month | 308868.5 | <0.0001 | -0.77 | Large |
| Precipitation Seasonality | 5884104 | <0.0001 | 0.86 | Large |
| Precipitation of Wettest Quarter | 5759096.5 | <0.0001 | 0.824 | Large |
| Precipitation of Driest Quarter | 316006.5 | <0.0001 | -0.768 | Large |
| Precipitation of Warmest Quarter | 5867988.5 | <0.0001 | 0.855 | Large |
| Precipitation of Coldest Quarter | 299375 | <0.0001 | -0.772 | Large |
| Windspeed | 3972914.5 | <0.0001 | 0.302 | Moderate |

$r_{rb}$ = rank-biserial correlation coefficient

Very Small = < 0.1, Small = 0.1-0.3, Moderate = 0.3-0.5, & Large > 0.5

Supplemental Table B: Descriptive statistics of orb-weaving spiders observed during surveys.

| Species | Frequency of Location Present | Average: When Present | Frequency without Jorōs | Average: without Jorōs | Frequency with Jorōs | Average: with Jorō | Total Individuals Observed |
|---|---|---|---|---|---|---|---|
| *Trichonephila clavata* | 53 | 10.2 ± 1.8 | 0 | 0.0 | 53 | 10.2 ± 1.8 | 543 |
| *Micrathena mitrata* | 49 | 6.5 ± 0.85 | 27 | 6.7 ± 1.3 | 22 | 6.2 ± 1.1 | 319 |
| *Neoscona crucifera* | 44 | 1.7 ± 0.17 | 21 | 2.1 ± 0.31 | 23 | 1.4 ± 0.15 | 76 |
| *Araneus marmoreus* | 27 | 1.3 ± 0.10 | 14 | 1.1 ± 0.10 | 13 | 1.4 ± 0.18 | 34 |
| *Verrucosa arenata* | 25 | 1.4 ± 0.10 | 13 | 1.5 ± 0.14 | 12 | 1.4 ± 0.15 | 36 |
| *Leucauge venusta* | 24 | 2.8 ± 0.73 | 13 | 3.2 ± 1.3 | 11 | 2.5 ± 0.53 | 68 |
| *Neoscona domiciliorum* | 20 | 1.9 ± 0.29 | 11 | 2.0 ± 0.41 | 9 | 1.7 ± 0.44 | 37 |
| *Metepeira labyrinthea* | 14 | 2.4 ± 0.72 | 5 | 1.4 ± 0.25 | 9 | 3.0 ± 1.1 | 34 |
| *Mangora maculata* | 9 | 2.0 ± 0.53 | 6 | 1.5 ± 0.22 | 3 | 3.0 ± 1.5 | 18 |
| *Micrathena gracilis* | 8 | 1.3 ± 0.25 | 4 | 1.5 ± 0.50 | 4 | 1.0 ± 0.0 | 10 |
| *Gasteracantha cancriformis* | 8 | 2.6 ± 0.73 | 6 | 3.0 ± 0.93 | 2 | 1.5 ± 0.50 | 21 |
| *Trichonephila clavipes* | 7 | 1.1 ± 0.14 | 6 | 1.2 ± 0.17 | 1 | 1.0 ± NaC | 8 |
| *Cyclosa turbinata* | 6 | 1.3 ± 0.33 | 3 | 1.0 ± 0.0 | 3 | 1.7 ± 0.67 | 8 |
| *Micrathena sagittata* | 5 | 1.6 ± 0.25 | 2 | 1.5 ± 0.50 | 3 | 1.7 ± 0.33 | 8 |
| *Argiope aurantia* | 5 | 1.4 ± 0.40 | 1 | 1.0 ± NaC | 4 | 1.5 ± 0.50 | 7 |
| *Mecynogea lemniscata* | 3 | 3.3 ± 2.3 | 1 | 8 ± NaC | 2 | 1.0 ± 0.0 | 10 |
| Kleptoparasite | 3 | 1.3 ± 0.33 | 3 | 1.3 ± 0.33 | 0 | 0.0 | 4 |
| *Tetragnatha* sp. | 1 | 3.0 ± NaC | 1 | 3.0 ± NaC | 0 | 0.0 | 3 |
| *Mangora* sp. | 1 | 1.0 ± NaC | 1 | 1.0 ± NaC | 0 | 0.0 | 1 |
| *Uloborid* sp. | 1 | 3.0 ± NaC | 1 | 3.0 ± NaC | 0 | 0.0 | 3 |

Mean ± SE

NaC = unable to calculate

Supplemental Figure B: Mean abundance (with 95% CI) grouped by the presence/absence of *Trichonephila clavata* at the survey location.

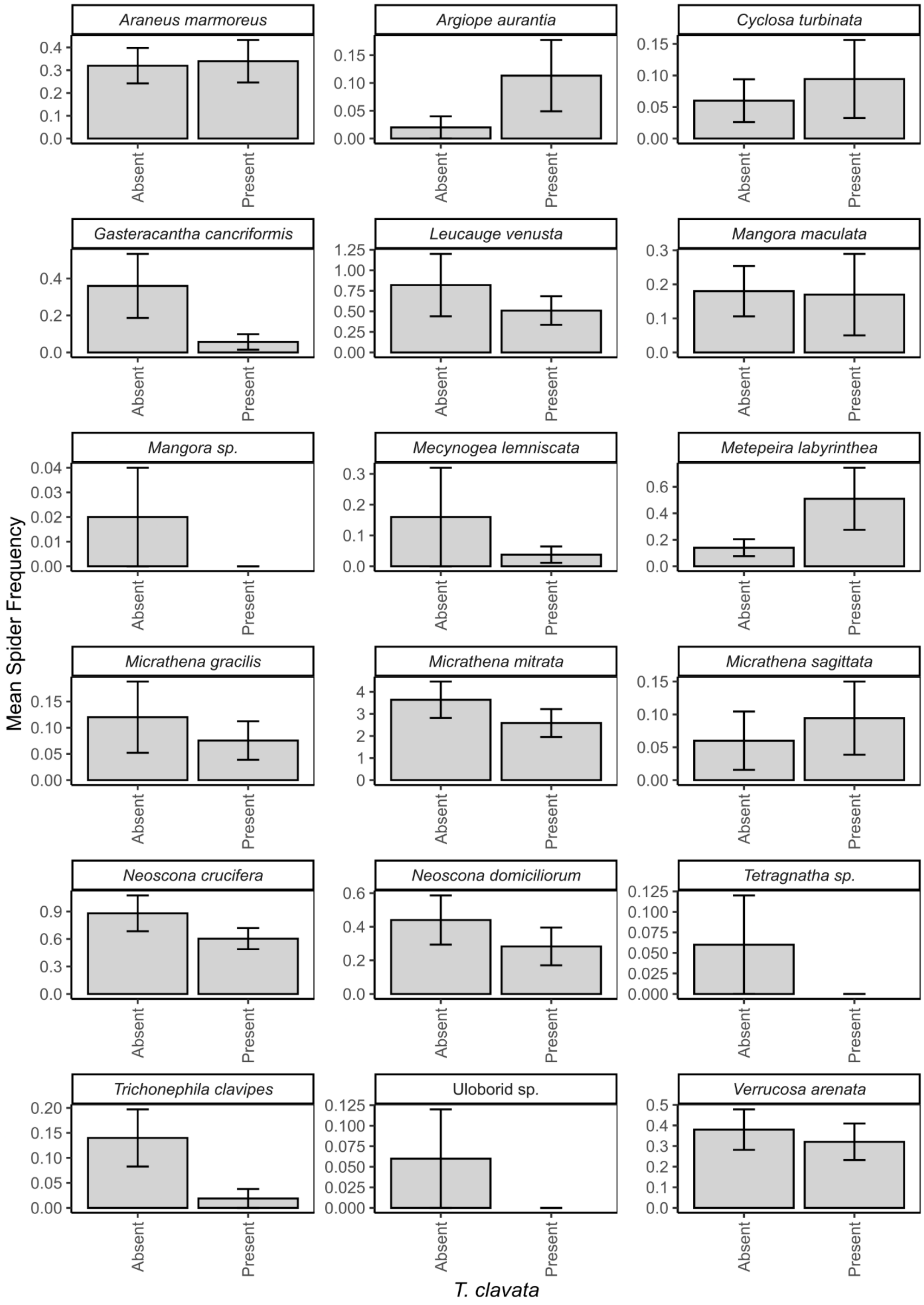

Supplemental Figure C: Interaction plots based on linear and generalized linear regression models of a location's human population density and a measure of *Trichonephila clavata's* presence (in space or time) on orb weaving spider diversity. Shading around trend lines represents 95% CI.

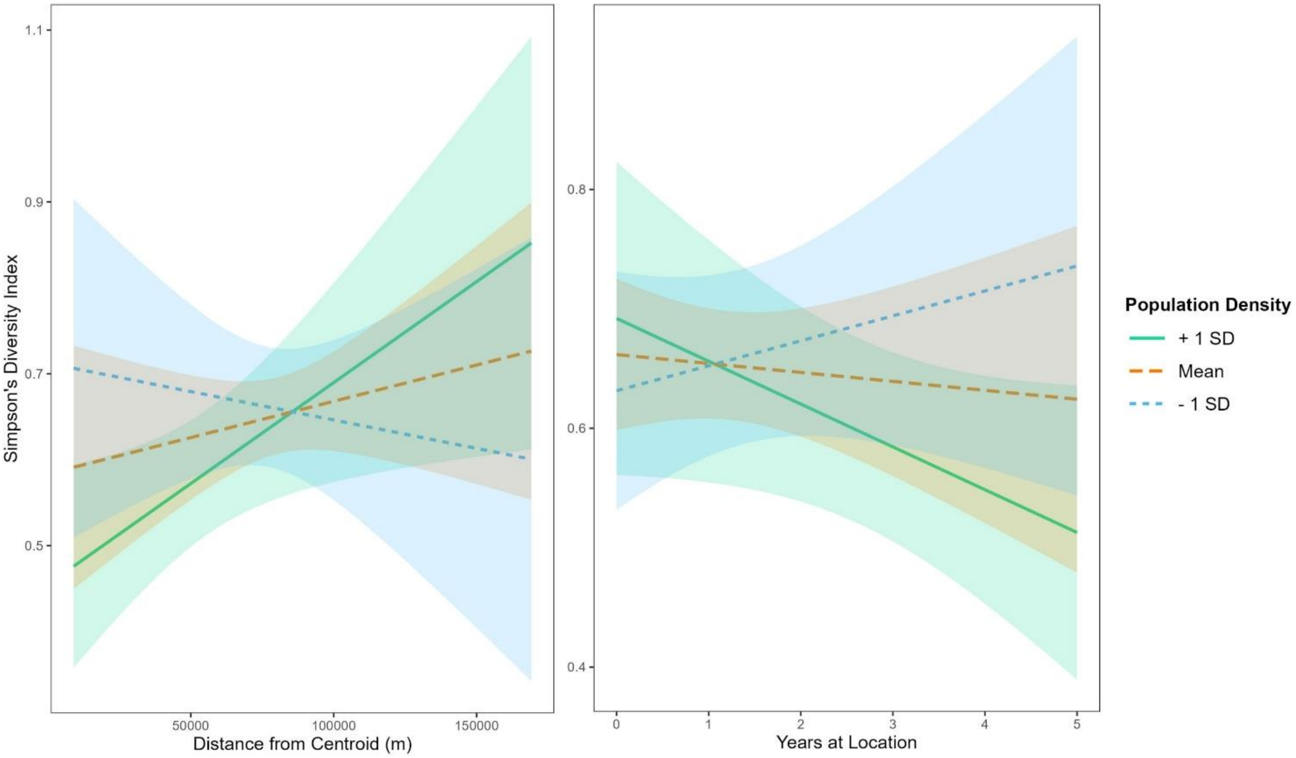
